# 3D architecture and structural flexibility revealed in the subfamily of large glutamate dehydrogenases by a mycobacterial enzyme

**DOI:** 10.1101/2020.11.14.381715

**Authors:** Melisa Lázaro, Roberto Melero, Charlotte Huet, Jorge P. López-Alonso, Sandra Delgado, Alexandra Dodu, Eduardo M. Bruch, Luciano A. Abriata, Pedro M. Alzari, Mikel Valle, María-Natalia Lisa

## Abstract

Glutamate dehydrogenases (GDHs) are widespread metabolic enzymes that play key roles in nitrogen homeostasis. Large glutamate dehydrogenases composed of 180 kDa subunits (L-GDHs_180_) contain long N- and C-terminal segments flanking the catalytic core. Despite the relevance of L-GDHs_180_ in bacterial physiology, the lack of structural data for these enzymes has limited the progress of functional studies. Here we show that the mycobacterial L-GDH_180_ (mL-GDH_180_) adopts a quaternary structure that is radically different from that of related low molecular weight enzymes. Intersubunit contacts in mL-GDH_180_ involve a C-terminal domain that we propose as a new fold and a flexible N-terminal segment comprising ACT-like and PAS-type domains that could act as metabolic sensors for allosteric regulation. These findings uncover unique aspects of the structure-function relationship in the subfamily of L-GDHs.

## Introduction

Glutamate dehydrogenases (GDHs) are ubiquitous oligomeric enzymes that catalyze the reversible oxidative deamination of L-glutamate to 2-oxoglutarate, at the crossroad between the Krebs cycle and ammonium assimilation. GDHs are grouped into the subfamily of small GDHs composed of subunits of 50 kDa (S-GDHs_50_) and the subfamily of large GDHs (L-GDHs) composed of monomers of 115 kDa (L-GDHs_115_) or 180 kDa (L-GDHs_180_) (Miñambres et al., 2000). L-GDHs, found in lower eukaryotes and prokaryotes, are NAD^+^ dependent enzymes that differ from S-GDHs_50_ by the presence of long N- and C-terminal extensions flanking the catalytic domain (Miñambres et al., 2000). The possible role(s) of such terminal segments in oligomerization and/or enzyme regulation has remained largely unknown (Beaufay et al., 2015; Camardella et al., 2002; Kawakami et al., 2007, 2010; Lu and Abdelal, 2001; Miñambres et al., 2000; Nott et al., 2009; O’Hare et al., 2008; Veronese et al., 1974).

The relevance of L-GDHs_180_ in bacterial physiology has been emphasized in previous studies of environmental (Beaufay et al., 2015) and pathogenic species (DeJesus et al., 2013; Griffin et al., 2011). Among the later, the mycobacterial L-GDH_180_ (mL-GDH_180_) is part of a signal transduction pathway that senses amino acid availability to control metabolism and virulence of *Mycobacterium tuberculosis* (Nott et al., 2009; Rieck et al., 2017; York, 2017). This enzyme is essential for the *in vitro* growth of the tubercle bacillus (DeJesus et al., 2013; Griffin et al., 2011) whereas it is crucial for *Mycobacterium bovis* BCG survival in media containing glutamate as the sole carbon source (Gallant et al., 2016). Moreover, diverse mechanisms have been implicated in the regulation of L-GDHs_180_. The catabolism of glutamate by mL-GDH_180_ is inhibited by the regulator GarA (Nott et al., 2009; O’Hare et al., 2008) when extracellular nitrogen donor amino acids are available (Rieck et al., 2017) whereas the L-GDH_180_ from *Streptomyces clavuligerus* (Miñambres et al., 2000) (filo Actinobacteria, which includes mycobacteria) as well as L-GDHs_180_ from Proteobacteria (Kawakami et al., 2007, 2010; Lu and Abdelal, 2001) are directly regulated by amino acids. Despite the key roles of L-GDHs_180_ in the redistribution of amino groups within cells, their 3D structure has remained elusive, preventing a deeper understanding of the molecular basis of enzyme function.

Here we report the 3D structure of the mL-GDH_180_ isoform from *Mycobacterium smegmatis*, obtained through an integrative approach that combined single-particle cryo-EM and X-ray protein crystallography data at resolutions between 4.11 and 6.27 Å. Our findings reveal unique characteristics of domain organization and oligomeric assembly in the L-GDHs subfamily, thus allowing to update the annotation of the Pfam family PF05088 that includes the L-GDHs_180_, and offer a rationale for the direct regulation of L-GDHs_180_ by metabolites. Furthermore, our cryo-EM data uncover fluctuations of the quaternary structure of mL-GDH^180^ that are possibly relevant for the allosteric regulation of the enzyme activity.

## Results

### The 3D architecture of mL-GDH_180_

As revealed by X-ray protein crystallography and single-particle cryo-EM (Figure 1 and Figure 2), mL-GDH_180_ assembles into a homotetramer. mL-GDH_180_ monomers are arranged around perpendicular two-fold axes that pass through a central cavity in the structure.

**Figure 1.**
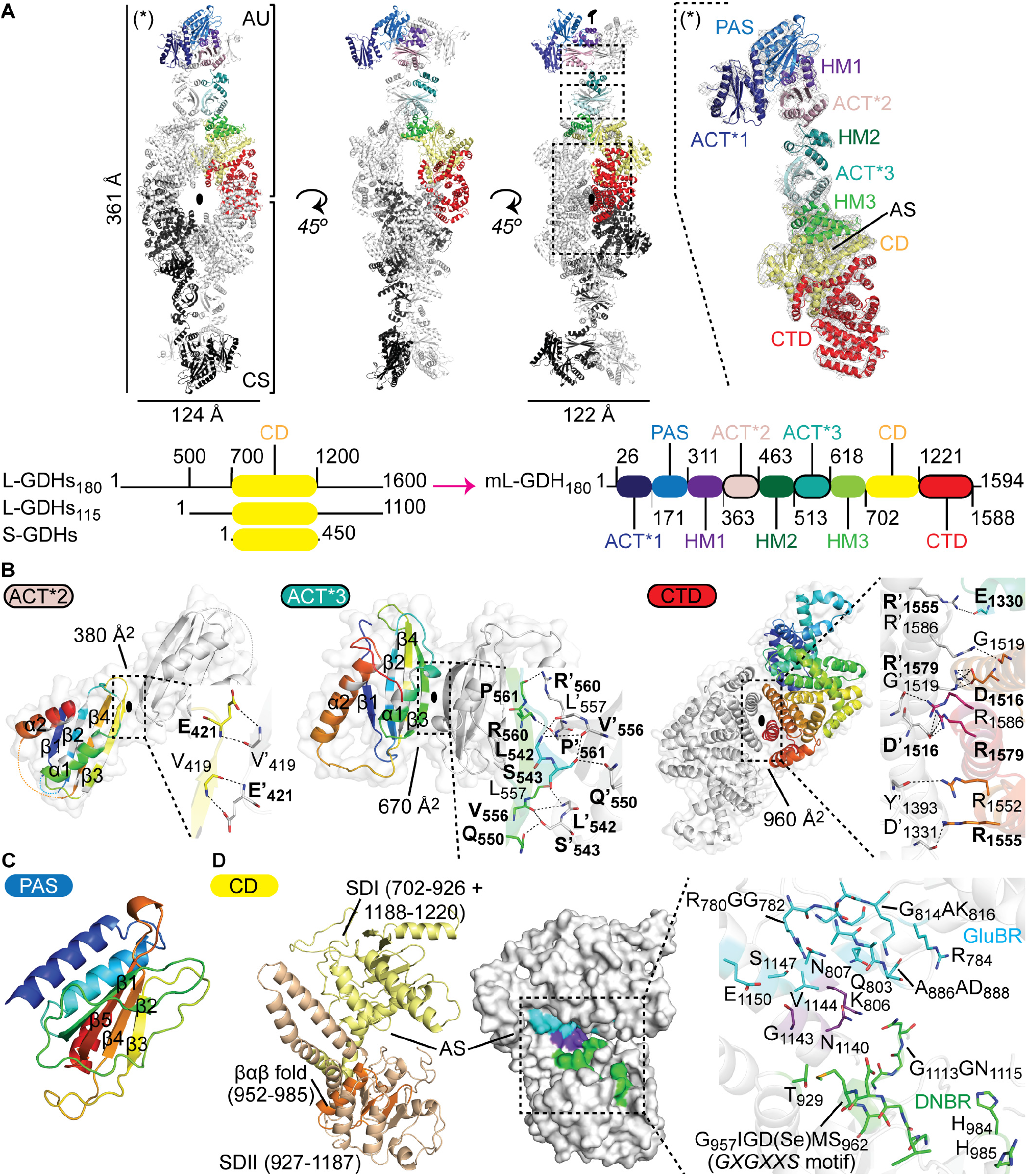
Crystal structure of Se-Met mL-GDH_180_. (**A**) The asymmetric unit (AU) contains two monomers (RMSD of 0.51 Å for 289 alpha carbons in segment 45-362, 0.26 Å for 1163 alpha carbons in segment 368-1588); a tetramer (as ribbons) is formed by crystallographic symmetry (CS); oval symbols represent two-fold axes. The *2mFo–DFc* electron density (gray mesh), contoured to 1.5 σ, is shown for one protein subunit on the right. Domains boundaries are given in residue numbers in a scheme below; CD, catalytic domain; CTD, C-terminal domain; AS, active site. A comparative scheme of L-GDHs180, L-GDHs115 and S-GDHs_50_ is also provided, with approximate residue numbers. (**B**) Oligomeric interfaces (areas in Å^2^) involve the domains ACT*2, ACT*3 and CTD. Contacting residues (as sticks in insets) labeled in bold characters are strictly conserved in diverse L-GDHs. The topology of domains ACT*2 and ACT*3 is highlighted with rainbow colors; white positions within the rainbow depict conserved core residues (Lang et al., 2014). (**C**) The PAS domain. (**D**) The CD is shown with the SDI and SDII in yellow and orange, respectively. The βαβ motif is involved in dinucleotide binding (Miñambres et al., 2000). The glutamate-binding region (GluBR, cyan) and the dinucleotide-binding region (DNBR, green) (Miñambres et al., 2000) are highlighted in a surface representation and as sticks in an inset. Residues in purple conform both binding regions (Miñambres et al., 2000).

**Figure 2.**
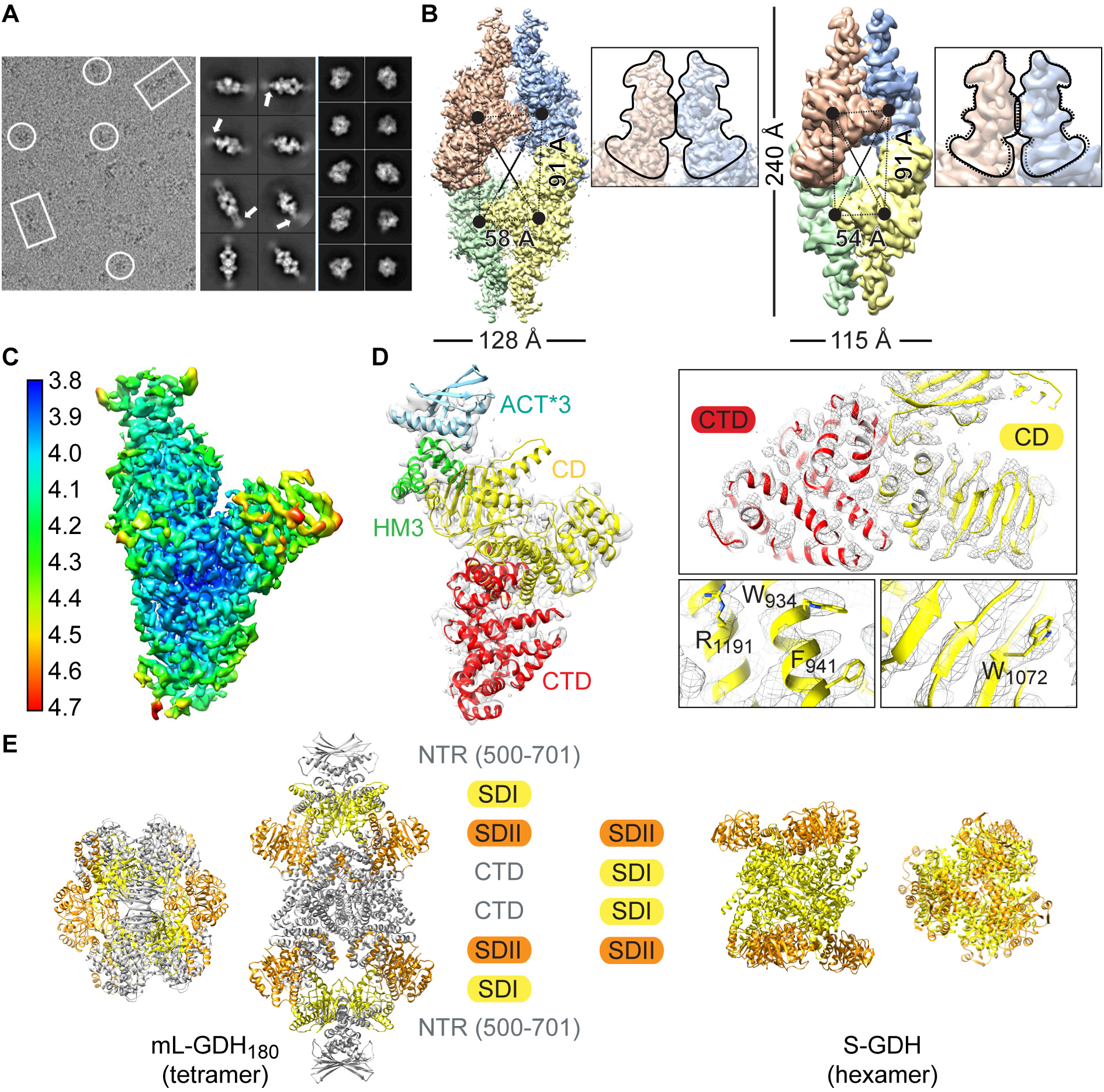
Intrinsic flexibility and alternate conformers of native mL-GDH_180_. (**A**) Cryo-EM image obtained for mL-GDH_180_ (left panel) showing side (rectangles) and top (circle) views for single particles. The 2D class averages for mL-GDH_180_ tetramers (right panels) display flexible ends at side views (white arrows). (**B**) Cryo-EM density maps for the open (left, 4.47 Å resolution) and close (right, 6.6 Å resolution) conformations of mL-GDH_180_ tetramers, segmented into the four subunits. (•): centers of mass of the subunits. Insets are close-up views of the contact zone between the N-terminal regions (NTRs, contoured as —) of two monomers; the contour of the NTRs of the open form is also shown as (---) on the closed conformation, for comparison. (**C**) Local resolution for a single subunit of the open conformation after focused refinement (average resolution is 4.11 Å). (**D**) Cryo-EM map for one mL-GDH^180^ subunit and the fitted atomic coordinates. Domains colors and labels are as in Figure 1. Insets are close-up views; selected amino acid side chains are shown as sticks. (**E**) Comparison of the quaternary structure of mL-GDH_180_ and a representative hexameric S-GDH_50_ (PDB code 3SBO). The catalytic domains are colored into SDI (yellow) and SDII (orange). The NTRs (only the portion that is well defined in cryo-EM maps is displayed) and the CTDs of mL-GDH_180_ monomers are depicted in gray.

The 6.27 Å resolution crystal structure of the seleno-methionine (Se-Met) derivative of mL-GDH_180_ (Figure 1 and Table 1), obtained as illustrated in Figure S1 through an integrative strategy that also included cryo-EM data up to 4.11 Å, revealed that the protein subunits display a unique domain organization (Figure 1A). The N-terminal segment comprises three ACT (*A*spartate kinase-*C*horismate mutase-*T*yrA) −like (Lang et al., 2014) (hereafter ACT*, see below) domains (ACT*1-3), a PAS (*P*er-*A*rnt-*S*im) −type (Möglich et al., 2009) domain and three helical motifs (HM1-3). Notably, the primary structures of ACT and PAS domains are poorly conserved and, therefore, these modules are often difficult to identify from BLAST searches (Lang et al., 2014; Möglich et al., 2009). The C-terminal region consists of a single helical domain that showed no detectable structural similarity to previously characterized proteins in Dali (Holm, 2020), ECOD (Cheng et al., 2014), CATH (Dawson et al., 2017) and VAST (Madej et al., 2014) searches and, therefore, constitutes a possible new fold.

**Table 1.**
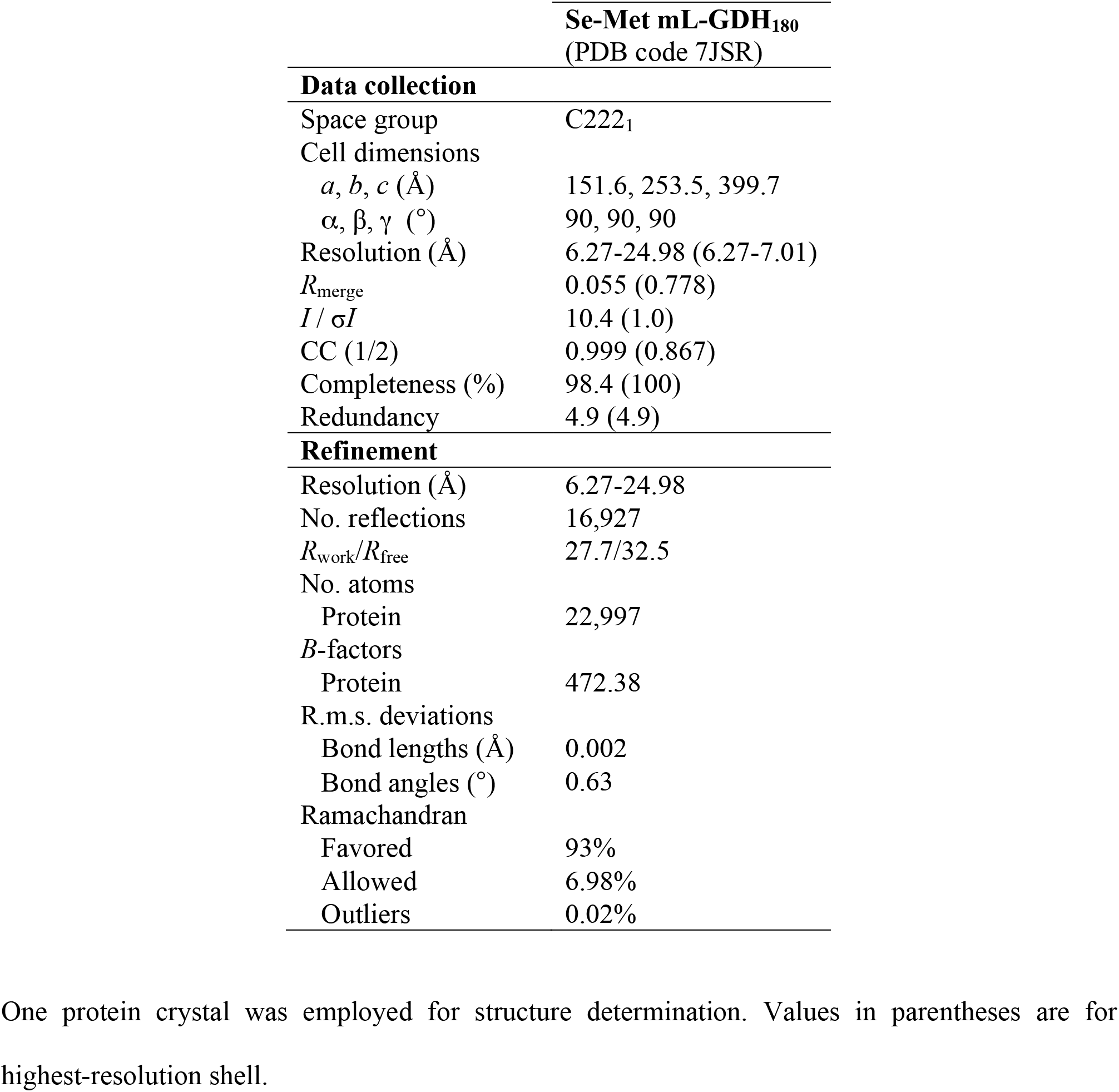
X-ray diffraction data collection and refinement statistics.

The catalytic domains in the mL-GDH_180_ complex were not found to contribute intersubunit contacts (Figure 1A). Instead, the N- and C-terminal regions of mL-GDH_180_ provide dimer-like interactions between pairs of monomers. Contacts between mL-GDH_180_ subunits engage the ACT*2, ACT*3 and C-terminal domains (Figure 1B). Most of the residues involved in interfacial hydrogen bonds or salt bridges in mL-GDH_180_ are strictly conserved in the enzyme isoform from *M. tuberculosis* (O53203, 72% sequence identity) (Nott et al., 2009), the L-GDH_180_ from *S. clavuligerus* (E2Q5C0, 47% sequence identity) (Miñambres et al., 2000) and the L-GDH_115_ from *Nocardia farcinica* (A0A0H5NTF9, 55% sequence identity over non-gap aligned columns). Except for a single amino acid (Arg560), the same group of residues is also conserved in the L-GDH_180_ from *P. aeruginosa* (Q9HZE0, 40% sequence identity) (Lu and Abdelal, 2001). These observations underscore the functional relevance of the oligomeric assembly found for mL-GDH_180_.

ACT and PAS modules are known to regulate functionally diverse proteins by driving conformational and/or quaternary structural changes (Lang et al., 2014; Möglich et al., 2009). The binding of specific amino acids to ACT-ACT interfaces confers allosteric control to oligomeric enzymes involved in amino acid metabolism (Lang et al., 2014) whereas PAS modules sense and transduce chemical or physical stimuli to typically dimeric effector domains (Möglich et al., 2009). The ACT* domains of mL-GDH_180_ differ from the archetypal ACT fold in that strand β_1_ is located in the position usually occupied by strand β_4_, creating an ACT-like ββαββα topology with a β_1_β_2_β_4_β_3_ antiparallel sheet (Figure 1B and Figure S2). Similar variations of the characteristic ACT fold have been described for aspartate kinases and a mammalian tyrosine hydroxylase (Lang et al., 2014; Zhang et al., 2014), including sixteen core residues that are conserved in the ACT*1-3 domains of mL-GDH_180_ (Figure S2). Notably, the interaction between ACT*3 modules in mL-GDH_180_ produces a continuous eight-stranded antiparallel β-sheet with helices on one side (Figure 1B). A similar side-by-side arrangement of ACT domains generates allosteric amino acid binding sites in 3-phosphoglycerate synthases and aspartate kinases (Lang et al., 2014). Close to a dimer-like interface, the PAS module in mL-GDH_180_ adopts a typical fold (Figure 1C), comprising a core five-stranded β-sheet usually involved in signal sensing (Möglich et al., 2009), and displays up to 12% sequence identity with PAS domains in sensor histidine kinases retrieved in Dali (Holm, 2020) searches.

Similarly to S-GDHs_50_, the catalytic core of mL-GDH_180_ consists of subdomains SDI and SDII (Figure 1D), with the active site located in a groove in-between. Functionally important residues in the catalytic domain of L-GDHs_180_ have been previously identified by their conservation in sequence comparisons of diverse GDHs (Miñambres et al., 2000). The SDI in mL-GDH_180_ contains most of the residues of the glutamate-binding region whereas the SDII conforms the dinucleotide-binding site.

### Intrinsic flexibility and alternate conformers of mL-GDH_180_

Cryo-EM and SAXS data uncovered the intrinsic flexibility of native mL-GDH_180_ (Figure 2, Figure S3 and Table S1). 2D averages for side views of mL-GDH_180_ tetramers revealed a high degree of flexibility at distal ends, where ACT*1-2 and PAS domains reside, and their corresponding densities vanished in 3D cryo-EM maps (Figure 2A). A 3D-classification of the detected mL-GDH^180^ particles was performed to distinguish alternate conformers of the enzyme. Two mL-GDH_180_ conformers were found, called the open and close conformations (Figure 2B), for which the ACT*3 module, the HM3, the catalytic domain and the C-terminal region were defined in each monomer, achieving an estimated 4.11 Å resolution for this region in the open conformation (Figure 2C-D, Figure S4 and Table 2). The two conformers differ in the relative positions of the centers of mass of the subunits (Figure 2B). The catalytic domains in mL-GDH_180_ monomers in contact through their N-terminal segments are found closer to each other in the less stable close conformation compared to the open form. Overall, these findings reveal transitions of the quaternary structure that could intervene in the allosteric regulation of the enzyme.

**Table 2.** Cryo-EM data collection and processing.

|  | **Open form**<br>(EMD-11606) | **Close form**<br>(EMD-11612) | **Monomer**<br>(EMD-11613) |
| --- | --- | --- | --- |
| Magnification | 47,170 | 47,170 | 47,170 |
| Voltage (kV) | 300 | 300 | 300 |
| Electron dose ($e^-/\text{\AA}^2$) | 28 (14 fractions) | 28 (14 fractions) | 28 (14 fractions) |
| Defocus range ($\mu\text{m}$) | 0.67-3.26 | 0.67-3.26 | 0.67-3.26 |
| Pixel size ($\text{\AA}$) | 1.06 | 1.06 | 1.06 |
| Symmetry imposed | D2 | D2 | C1 |
| Initial particles images | 276,704 | 276,704 | 276,704 |
| Final particles images | 63,715 | 42,476 | 63,715 x 4 |
| Map resolution ($\text{\AA}$) | 4.47 | 6.6 | 4.11 |
| Map sharpening B factor | 250 | 250 | 250 |

## Discussion

L-GDHs_180_ were discovered in 2000 from a study of *Streptomyces clavurigerus* (Miñambres et al., 2000) and were later isolated from other diverse bacterial species, including *Pseudomonas aeruginosa* (Lu and Abdelal, 2001), psychrophilic bacteria (Camardella et al., 2002; Kawakami et al., 2007, 2010), *Caulobacter crescentus* (Beaufay et al., 2015) and *Mycobacterium* spp (Nott et al., 2009; O’Hare et al., 2008). As sequences of L-GDHs were identified, they were analyzed in light of the available crystallographic evidence for S-GDHs_50_ (Britton et al., 1992; Miñambres et al., 2000). S-GDHs_50_ are hexameric enzymes in which the oligomeric interfaces are conformed by motifs that are located within the catalytic domain (Britton et al., 1992; Miñambres et al., 2000). Most of these motifs are substantially modified in L-GDHs, either through sequence changes, insertions or deletions (Britton et al., 1992; Miñambres et al., 2000). In agreement with proposals that the oligomeric assembly would then be different for the two enzyme subfamilies (Britton et al., 1992; Miñambres et al., 2000), the quaternary structure of mL-GDH_180_ depends on interactions established by the N- and C-terminal regions flanking the catalytic domain (Figure 1A) and is radically different from that of S-GDHs_50_ (Figure 2E). The stoichiometry of the mL-GDH^180^ complex observed by cryo-EM and X-ray protein crystallography (Figure 1 and Figure 2) is supported by molecular weight estimates from SAXS data (Table S1) and is consistent with previous reports of tetrameric complexes of L-GDHs studied in solution (Lu and Abdelal, 2001; Veronese et al., 1974). Furthermore, most of the residues involved in interactions between mL-GDH_180_ monomers are conserved (Figure 1B) not only in mycobacterial isoforms of the enzyme but also in L-GDHs from diverse species in Actinobacteria and Proteobacteria. This suggests that the oligomeric assembly of mL-GDH_180_ may be a common theme in the enzyme subfamily.

The catalytic domains in the mL-GDH_180_ complex are oriented opposite to those in S-GDHs_50_ (Figure 2E), with the SDI (Figure 1D) directed toward the distal ends of the protein, where the monomers N-terminal region resides. This segment comprises ACT-like modules as well as a PAS-like domain arranged in tandem (Figure 1A) and shows a high degree of flexibility (Figure 2A). A comparison of the mL-GDH_180_ conformers identified by cryo-EM (Figure 2B) shows that conformational changes in the N-terminal region correlate with alterations in the relative positions of the catalytic domains. Taking into account the known roles of ACT modules in the allosteric control of oligomeric enzymes involved in amino acid metabolism (Lang et al., 2014), our findings offer a rationale for previous evidence pointing out the direct regulation of diverse L-GDHs_180_ by metabolites (Lu and Abdelal, 2001; Miñambres et al., 2000).

In conclusion, our findings suggest that the N-terminal segment of mL-GDH_180_ (as well as in related enzymes) could transduce intracellular metabolic stimuli to the catalytic core by driving changes in the quaternary structure. The reported 3D model of mL-GDH_180_ can now frame future studies to dissect the structure-function relationship of this enzyme and other members of the L-GDHs subfamily.

## Acknowledgments

The authors acknowledge the support and the use of resources of Instruct, a Landmark ESFRI project and Instruct Image Processing Center (I2PC) through Instruct Access Project PID6258, NVIDIA Corporation for the donation of the Quadro GP100 GPU used for this research, the eBIC Electron Bio-Imaging Centre (Diamond light source, Oxford) and the Netherlands Centre for Electron Nanoscopy (Leiden, Netherlands) for the collection of cryo-EM images, and the European Synchrotron Radiation Facility (Grenoble, France) for the collection of diffraction and SAXS data. We thank David Gil, Ariel Mechaly, Ahmed Haouz and Pascal Le Normand for technical advice and insightful discussions. This work was supported by the grant PICT 2017-1932, from the Agencia Nacional de Promoción de la Investigación, el Desarrollo Tecnológico y la Innovación (Agencia I+D+i, Argentina), received by MNL, and the grant PGC2018-098996-B-100 from the Spanish Ministerio de Ciencia e Innovación, received by MV. MV thanks the AEI (Agencia Estatal de Investigación) for the Severo Ochoa Excellence Accreditation (SEV-2016-0644).

## Author contributions

ML produced protein, prepared EM samples, performed EM experiments, processed and analysed EM data, and performed structural analyses; RM determined the initial cryo-EM model; CH optimized protein production; JPLA processed and analysed EM data; SD contributed to protein production and the preparation of EM samples; AD contributed to protein production; EMB analysed SAXS data; LAA contributed to *ab initio* modelling and structural analyses; PMA designed experiments and analysed data; MV designed experiments, attended EM data collection and processing and analysed results; MNL cloned the gene of mL-GDH_180_, optimized protein production, produced protein, obtained protein crystals, solved the crystal structure of the protein, refined the structure of mL-GDH_180_ obtained by cryo-EM, designed experiments, acquired data, analysed data and wrote the paper. All authors read and corrected the paper.

## Deposition of structures and maps

Cryo-EM maps obtained for mL-GDH_180_ were deposited in the Electron Microscopy Data Bank under the accession codes EMD-11606 (open conformation), EMD-11612 (close conformation) and EMD-11613 (monomer). Atomic coordinates for the open form of mL-GDH_180_ derived from cryo-EM data were deposited in the Protein Data Bank under the accession code 7A1D. Structure factors and atomic coordinates obtained for Se-Met mL-GDH_180_ by X-ray protein crystallography were deposited in the Protein Data Bank under the accession code 7JSR.

## Methods

### Protein production and purification

The sequence coding for the L-GDH_180_ from *M. smegmatis* MC^2^-155 (MSMEG_4699, Uniprot A0R1C2) was cloned into vector pLIC-His (Cabrita et al., 2006) employing the oligonucleotides Fw: CCAGGGAGCAGCCTCGATGATTCGCCGGCTTTCGG and Rv: GCAAAGCACCGGCCTCGTTACCCAGTCGTTCCGGTCCC. The resulting plasmid was used to produce N-terminally His6-tagged mL-GDH_180_ in *E. coli* cells. Transformed *E. coli* cells were grown at 37°C in medium supplemented with ampicillin or carbenicillin until reaching 0.8 units of optical density at 600 nm. Protein expression was then induced by adding isopropyl β-D-1-thiogalactopyranoside (IPTG) to a final concentration of 0.5 mM, and the incubation was continued for 18 hours at 14°C. Cells were harvested by centrifugation and sonicated. Following clarification by centrifugation, the supernatant was loaded onto a HisTrap HP column (GE Healthcare) equilibrated with buffer 25 mM HEPES, 500 mM NaCl, 20% v/v glycerol, 20 mM imidazole, pH 8.0, and His6-tagged mL-GDH_180_ was purified by applying a linear imidazole gradient (20-500 mM). The protein was then further purified by size-exclusion chromatography, as described bellow. mL-GDH_180_ containing fractions, as confirmed by SDS-PAGE and measurements of glutamate dehydrogenase activity (O’Hare et al., 2008), were pooled and used immediately. The protein was quantified by electronic absorption using the molar absorption coefficient of 171,090 M^-1^ cm^-1^, predicted from the amino acid sequence by the ProtParam tool (http://web.expasy.org/protparam/). For EM and SAXS experiments, native mL-GDH_180_ was produced in *E. coli* BL21(DE3) cells grown in LB broth. Size-exclusion chromatography was performed using a Superose 6 10/300 GL column (GE Healthcare) equilibrated in buffer 20 mM MES, 300 mM NaCl, 5 mM MgCl_2_, pH 6.0. Instead, Se-Met mL-GDH_180_ for crystallographic studies was produced in *E. coli* B834 (DE3) cells grown in SelenoMethionine Medium Complete (Molecular Dimensions), and size-exclusion chromatography was carried out using a HiPrep Sephacryl S-400 HR column (GE Healthcare) equilibrated in buffer 25 mM Tris, 150 mM NaCl, pH 7.5.

GarA from *M. tuberculosis* was produced as previously described (England et al., 2009).

### Cryo-electron microscopy

4 μl of 0.3 mg/ml mL-GDH_180_ were applied to Quantifoil R2/2 holey carbon grids and vitrified using a Vitrobot (FEI). Data collection was carried out in a Titan Krios FEI electron microscope operated at 300 kV by a K2 direct detector (GATAN). Movie frames (1,802) were taken at a nominal magnification of x 47,170 resulting in a sampling of 1.06 Å/pixel. Each movie contained 20 frames with an accumulated dose of 40 *e*^-^/Å^2^. Movie frames were aligned using MotionCor (Li et al., 2013; Zheng et al., 2017), and the final average included frames 2-15 with a total dose of 28 *e*^-^/Å^2^ on the sample.

The contrast transfer function (CTF) of the micrographs was estimated using CTFFIND4 (Rohou and Grigorieff, 2015). The particles were automatically selected from the micrographs using autopicking from RELION-3 (Zivanov et al., 2018). Evaluation of the quality of particles and selection was performed after 2D classifications with SCIPION (de la Rosa-Trevín et al., 2016) and RELION-3 (Zivanov et al., 2018) software packages. The initial volume for 3D image processing was calculated using common lines in EMAN (Tang et al., 2007) and using the algorithm 3D-RANSAC (Vargas et al., 2014). With this initial reference, additional rounds of automated particle picking were performed. An initial data set of 276,704 particles was subjected to 2D and 3D class averaging in order to select the best particles. The 3D-classification of the 106,190 final particles with imposed D2 symmetry resulted in two different conformations, a close (40%) and an open form (60%), with estimated resolutions of 6.6 Å and 4.47 Å, respectively. To improve the blurred regions of the cryo-EM maps the refinement was focused on the subunits and the final resolution was 4.11 Å for a monomer in the open conformation. This refinement focused on single mL-GDH_180_ subunits was performed after the alignment of all the monomers following the D2 symmetry, with masked subunits. Local resolution was estimated using RELION-3 (Kucukelbir et al., 2014; Zivanov et al., 2018).

Model fitting into cryo-EM maps was performed using the programs UCSF Chimera (Pettersen et al., 2004), Namdinator (Kidmose et al., 2019), phenix.real_space_refine (Afonine et al., 2018) and Coot (Emsley et al., 2010). Residues 500-1588 from the crystal structure of Se-Met mL-GDH_180_ (see below) were fitted into the cryo-EM map of the open form of the protein. Se-methionine residues were replaced by methionine residues using Coot (Emsley et al., 2010) and the model was finally refined employing phenix.real_space_refine (Afonine et al., 2018) with NCS and secondary structure restraints. Overall correlation coefficients were: CC (mask): 0.73; CC (volume): 0.71; CC (peaks): 0.61. The final model contained 92% of the residues within favored regions of the Ramachandran plot and no outliers.

Figures were generated and rendered with UCSF Chimera (Pettersen et al., 2004).

Cryo-EM maps obtained for mL-GDH_180_ were deposited in the Electron Microscopy Data Bank under the accession codes EMD-11606 (open conformation), EMD-11612 (close conformation) and EMD-11613 (monomer). Atomic coordinates for the open form of mL-GDH^180^ derived from cryo-EM data were deposited in the Protein Data Bank under the accession code 7A1D.

### Negative staining electron microscopy

Negative-stained grids of mL-GDH_180_ were prepared using 2% uranyl acetate and visualized on a JEM-1230 transmission electron microscope (JEOL Europe) at an acceleration voltage of 80 kV. Images were taken in low dose conditions at a nominal magnification of x 30,000 using a GATAN CCD camera, resulting in 2.3 Å/pixel sampling.

Labeling of N-terminally His6-tagged mL-GDH_180_ was performed by direct incubation of electron microscopy grids in solutions containing 5 nm Ni-NTA-Nanogold (Nanoprobes). Briefly, after glow discharging the grids, the protein was incubated for 1 minute on the grids, fixed with 2% paraformaldehyde for 10 minutes at 4°C, washed 5 minutes with PBS, incubated for 15 minutes with Nanogold diluted 1/75 in PBS, washed twice with PBS, and finally stained with 2% uranyl acetate for 45 seconds.

### Crystallization, X-ray data collection and structure determination

Crystallization screenings were carried out using the sitting-drop vapor diffusion method and a Mosquito (TTP Labtech) nanoliter-dispensing crystallization robot. Crystals of Se-Met mL-GDH_180_ grew after 4-6 months from a 16.5 mg/ml protein solution containing an equimolar amount of GarA from *M. tuberculosis*, by mixing equal volumes of protein solution and mother liquor (100 mM sodium cacodylate pH 5.8, 12% v/v glycerol, 1.25 M (NH_4_)_2_SO_4_), at 4 °C. Single crystals were cryoprotected in mother liquor containing 32% v/v glycerol and flash-frozen in liquid nitrogen. X-ray diffraction data were collected at the synchrotron beamline ID23-1 (European Synchrotron Radiation Facility, Grenoble, France), at 100 K, using wavelength 0.99187 Å. Diffraction data were processed using XDS (Kabsch, 2010) and scaled with Aimless (Evans and Murshudov, 2013) from the CCP4 program suite (Winn et al., 2011).

The crystal structure of Se-Met mL-GDH_180_ was solved by molecular replacement using the program Phaser (McCoy et al., 2007). As search probe we used the atomic coordinates of a model built as follows. First, a poly-Ala model of mL-GDH_180_ was obtained from a preliminary *ca*. 7 Å resolution cryo-EM map of the protein, by employing the program phenix.map_to_model (Terwilliger et al., 2018). Features of the catalytic domain in mL-GDH_180_ monomers became apparent in the model, suggesting that the N-terminus of the polypeptide chains was located at the tips of the particle. This was confirmed by labeling N-terminally His6-tagged mL-GDH_180_ with Ni-NTA-Nanogold (Nanoprobes) and visualizing particles by negative staining electron microscopy. Then, the catalytic domain of mL-GDH_180_ (residues 702-1220) was homology-modeled by using the structure of the S-GDH_50_ from *C. glutamicum* (PDB code 5GUD) as template and employing MODELLER (Sali and Blundell, 1994) as implemented in the HHpred server (Zimmermann et al., 2018). One copy of the model of the catalytic domain was rigid-body fitted into the 7 Å cryo-EM map of mL-GDH180, which allowed updating the starting poly-Ala model by correcting helical elements and incorporating strands corresponding to the catalytic domain in one monomer of mL-GDH_180_. From this, the D2 tetramer was then rebuilt by applying NCS operators detected by phenix.find_ncs (Liebschner et al., 2019) and the model was refined against the 7 Å cryo-EM map using phenix.real_space_refine (Afonine et al., 2018) with NCS and secondary structure restraints. Finally, one of the protein chains in the resulting model was used as search probe to solve the crystal structure of Se-Met mL-GDH_180_ by molecular replacement.

Two monomers were placed within the asymmetric unit, which taken together with nearby crystallographic symmetry mates replicate the quaternary structure observed by cryo-EM. After crystallographic refinement using phenix.refine (Afonine et al., 2012; Headd et al., 2012) with NCS and secondary structure restraints, *mFo-DFc* and *2mFo-DFc* electron density maps displayed rod-shaped electron density peaks that remained un-modeled at this stage and that most likely corresponded to helices in the N-terminal region of mL-GDH_180_. Phase improvement by density modification with RESOLVE (Terwilliger et al., 2007) provided additional evidence in support of such elements. The N-terminal segment of mL-GDH_180_ (residues 1-701) was modeled *ab initio* using RaptorX (Wang et al., 2017; Xu, 2018), one of the top-ranking *ab initio* structure prediction methods according to recent CASP evaluations (Abriata et al., 2018, 2019). Raptor X works by initially estimating residue-residue contacts from residue coevolution patterns and uses the predicted contacts to drive model building; such technique has proven highly successful especially when integrated with experimental data (multiple examples overviewed in (Abriata and Dal Peraro, 2020)). The residue-residue contact map predicted by RaptorX and the models produced from it revealed that the N-terminal segment of mL-GDH_180_ comprises an array of contiguous domains, which were subsequently individually rigid-body fitted into the electron density maps. Similarly, the C-terminal domain of mL-GDH_180_ (residues 1221-1594) was modeled *ab initio* employing RaptorX (Wang et al., 2017; Xu, 2018) and used to correct and complete the crystallographic model. Finally, un-modeled or poorly modeled segments in the CD were manually built employing Coot (Emsley et al., 2010) from a 4.11 Å resolution cryo-EM map obtained for a monomer of mL-GDH_180_. The structure was then further refined by iterative cycles of manual model building with Coot (Emsley et al., 2010), used to apply stereochemical restraints, and crystallographic refinement of atomic coordinates and individual B-factors using phenix.refine (Afonine et al., 2012; Headd et al., 2012) with NCS and secondary structure restraints. The final model contained 93% of the residues within favored regions of the Ramachandran plot and 0.2% of outliers. The crystallographic structure of Se-Met mL-GDH^180^ correctly explained the connecting loops and bulky amino acid side chains evidenced for residues 500-1588 by a 4.47 Å cryo-EM map of the protein. Furthermore, the position of Se-Met residues in the crystal structure of Se-Met mL-GDH_180_ matched the position of peaks in an anomalous difference map calculated with diffraction data acquired at 0.979338 Å (12.66 keV), the Se K-edge.

Even though Se-Met mL-GDH_180_ crystallized in the presence of GarA from *M. tuberculosis*, electron density maps did not reveal evidences of co-crystallization and molecular replacement attempts with Phaser (McCoy et al., 2007) using the atomic coordinates of GarA in PDBs 2KFU or 6I2P failed. The evidence of helical elements in all mL-GDH_180_ domains allows excluding the presence of GarA (an all beta protein) from modeled regions, particularly from those involved in crystal contacts (mL-GDH_180_ residues 1-500).

Figures were generated and rendered with UCSF Chimera (Pettersen et al., 2004) or Pymol version 1.8.x (Schrödinger, LLC).

Atomic coordinates and structure factors obtained for Se-Met mL-GDH_180_ were deposited in the Protein Data Bank under the accession code 7JSR.

### Small angle X-ray scattering

Synchrotron SAXS data were collected at BioSAXS ID14EH3 beamline (European Synchrotron Radiation Facility, Grenoble, France) and recorded at 15°C using a PILATUS 1M pixel detector (DECTRIS) at a sample-detector distance of 2.43 m and a wavelength of 0.931 Å, resulting momentum transfer (*s*) ranging from 0.009 to 0.6 Å^-1^.

mL-GDH_180_ was assayed at concentrations ranging from 1 to 14 mg/ml in buffer 25 mM Tris, 150 mM NaCl, pH 7.5. For the buffer and the samples, ten 2D scattering images were acquired and processed to obtain radially averaged 1D curves of normalized intensity versus scattering angle. In order to optimize background subtraction, buffer scattering profiles recorded before and after measuring every sample were averaged. Then, for each protein sample, the contribution of the buffer was subtracted. All subsequent data processing was performed using the ATSAS suite (Franke et al., 2017).

Average scattering curves corresponding to different protein concentrations were compared using PRIMUS (Franke et al., 2017; Konarev et al., 2003). To obtain the idealized scattering curve the low *s* region of the most diluted sample and the high *s* region of the most concentrated sample were merged. The values of the forward scattering intensity *I*(0), the radius of gyration *R*_g_ as well as the dimensionless Kratky plot were calculated using PRIMUS (Franke et al., 2017; Konarev et al., 2003). Guinier plots of independent average scattering curves evidenced a constant *R*_g_ at different protein concentrations. The Porod volume was estimated using DATPOROD (Franke et al., 2017) and an *s*_max_ value equal to 7.5/*R*_g_. The pairwise distance distribution function *p*(*r*) and the maximum particle dimension *D*_max_ were calculated using GNOM (Franke et al., 2017; Svergun, 1992) with a reduced χ^2^ value of 1.07 for curve fitting. After running DAMMIN (Franke et al., 2017; Svergun, 1999) the excluded volume was estimated as *V*_ex_=volume of a single dummy atom*number of dummy atoms/0.74). Finally, the MW was estimated from the Porod volume and the excluded volume.

