## Supplemental figures and tables for "3D architecture and structural flexibility revealed in the subfamily of large glutamate dehydrogenases by a mycobacterial enzyme"

### Supplemental Information

#### Figure S1

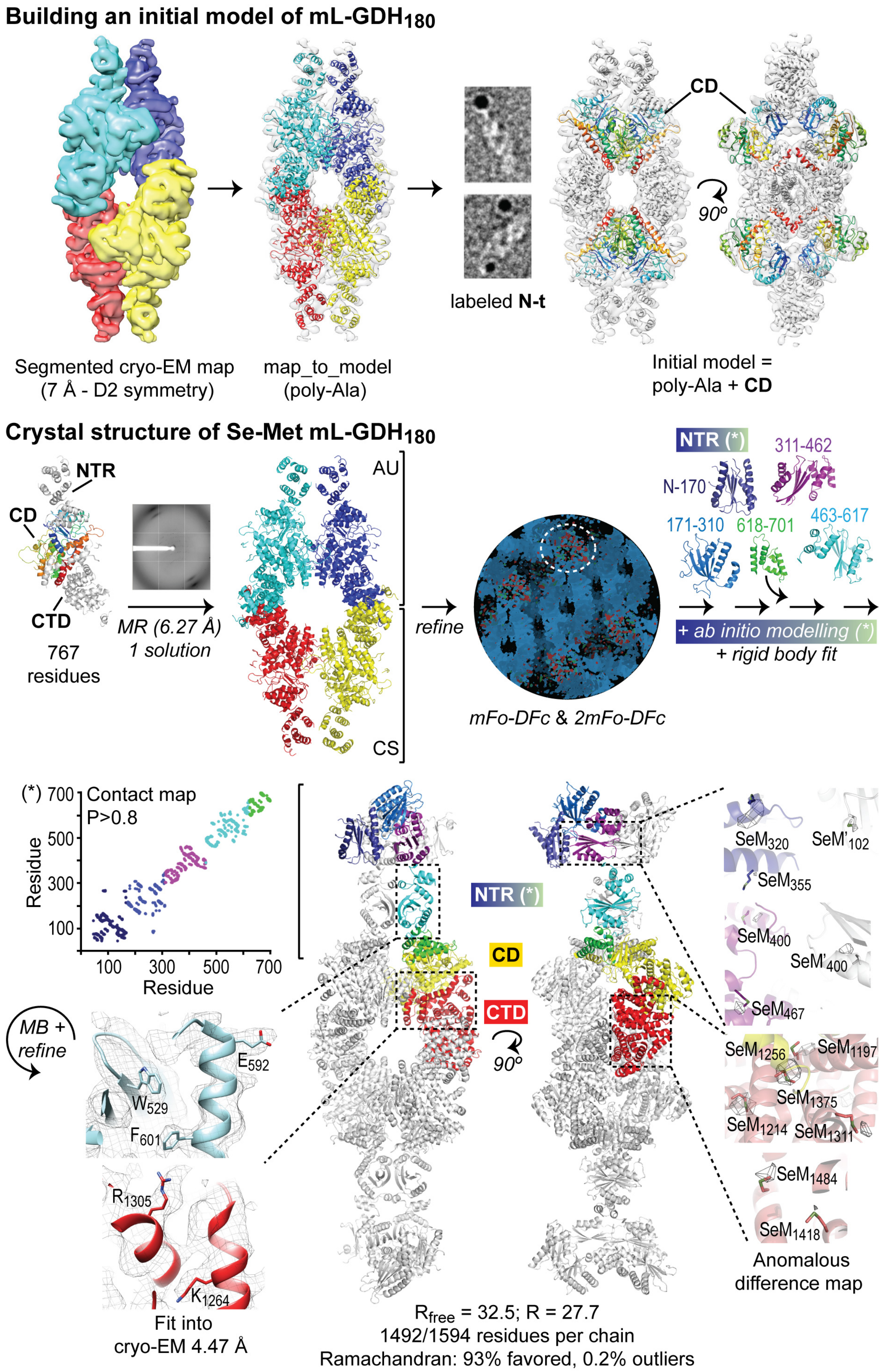

**Solving the crystal structure of Se-Met mL-GDH$_{180}$.** Upper panel: first, from a preliminary *ca.* 7 Å resolution cryo-EM map of mL-GDH$_{180}$ (segmented rendering at the left, with subunits in different colors; semi-transparent rendering at the right), we obtained a poly-Ala model (shown as ribbons) of the protein, by employing the program phenix.map_to_model (Terwilliger et al., 2018). Features of the catalytic domain in mL-GDH$_{180}$ monomers became apparent in the model, suggesting that the N-terminus (N-t) of the polypeptide chains was located at the tips of the particle. This was confirmed by labeling N-terminally His6-tagged mL-GDH$_{180}$ with Ni-NTA-Nanogold and visualizing particles by negative staining electron microscopy. Then, the catalytic domain of mL-GDH$_{180}$ (CD; residues 702-1220) was homology-modeled using the structure of the S-GDH$_{50}$ from *C. glutamicum* (PDB code 5GUD) as template and employing MODELLER (Sali and Blundell, 1994) as implemented in the HHpred server (Zimmermann et al., 2018). One copy of the model of the catalytic domain was rigid-body fitted into the 7 Å cryo-EM map of mL-GDH$_{180}$, which allowed updating the starting poly-Ala model by correcting helical elements and incorporating strands corresponding to the catalytic domain in one monomer of mL-GDH$_{180}$. From this, the D2 tetramer was then rebuilt by applying NCS operators detected by phenix.find_ncs (Liebschner et al., 2019) and the resulting model (poly-Ala + CD, with the CD in rainbow colors) was refined against the 7 Å cryo-EM map using phenix.real_space_refine (Afonine et al., 2018) with NCS and secondary structure restraints. Lower panel: we used one of the protein chains in the model poly-Ala + CD (767 residues) as search probe to solve the crystal structure of Se-Met mL-GDH$_{180}$ by molecular replacement with Phaser (McCoy et al., 2007). Two monomers were placed within the asymmetric unit (AU), which taken together with nearby crystallographic symmetry (CS) mates replicate the quaternary structure observed by cryo-EM. After crystallographic refinement using phenix.refine (Afonine et al., 2012; Headd et al., 2012) with NCS and secondary structure restraints, *mFo-DFc* and *2mFo-DFc* maps (in green and red contoured to 3 σ and in blue contoured to 1.5 σ, respectively) displayed rod-shaped electron density peaks that remained un-modeled at this stage (dashed circle) and that most likely corresponded to helices in the N-terminal region of mL-

GDH$_{180}$. Phase improvement by density modification with RESOLVE (Terwilliger et al., 2007) provided additional evidence in support of such elements. The N-terminal segment of mL-GDH$_{180}$ (NTS; residues 1-701) was modeled *ab initio* using RaptorX (Wang et al., 2017; Xu, 2018), one of the top-ranking *ab initio* structure prediction methods according to recent CASP evaluations (Abriata et al., 2018, 2019). RaptorX works by initially estimating residue-residue contacts from residue coevolution patters and uses the predicted contacts to drive model building; such technique has proven highly successful especially when integrated with experimental data (multiple examples overviewed in (Abriata and Dal Peraro, 2020)). The residue-residue contact map predicted by RaptorX ((*), showing contacts with probabilities (P) higher than 0.8) and the models produced from it (colored following the scheme of the contact map) revealed that the NTS comprises an array of contiguous domains, which were subsequently individually rigid-body fitted into the electron density maps. Similarly, the C-terminal domain of mL-GDH$_{180}$ (CTD; residues 1221-1594) was modeled *ab initio* employing RaptorX (Wang et al., 2017; Xu, 2018) and used to correct and complete the crystallographic model. Finally, un-modeled or poorly modeled segments in the CD were manually built employing Coot (Emsley et al., 2010) from a 4.11 Å resolution cryo-EM map obtained for a monomer of mL-GDH$_{180}$ (see Figure 2C-D and Figure S3). The structure was then further refined by iterative cycles of manual model building (MB) with Coot (Emsley et al., 2010), used to apply stereochemical restraints, and crystallographic refinement of atomic coordinates and individual B-factors using phenix.refine (Afonine et al., 2012; Headd et al., 2012) with NCS and secondary structure restraints. The final model contained 93% of the residues within favored regions of the Ramachandran plot and 0.2% of outliers (see also Table 1). The crystallographic structure of Se-Met mL-GDH$_{180}$ correctly explained the connecting loops (shown as ribbons) and bulky amino acid side chains (shown as sticks) evidenced for residues 500-1588 by a 4.47 Å cryo-EM map of the protein. Furthermore, the position of Se-Met residues (shown as sticks) in the crystal structure of Se-Met mL-GDH$_{180}$ matched the position of peaks in an anomalous difference map (shown as a gray mesh, contoured to 3 σ) calculated with diffraction data acquired at 0.979338 Å

(12.66 keV), the Se K-edge.

#### Figure S2

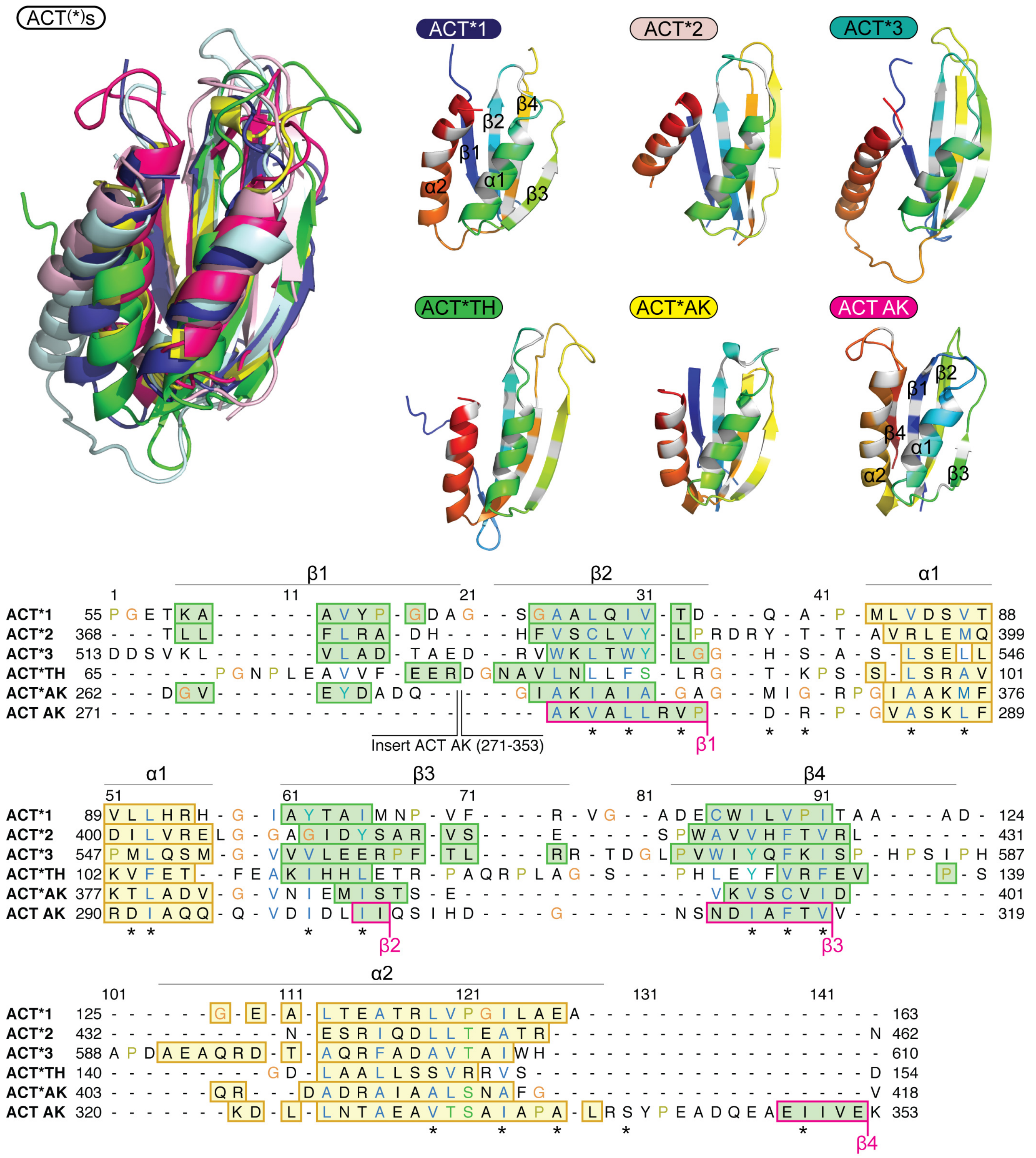


**Comparison of the ACT-like (ACT*) domains of mL-GDH$_{180}$.** Upper panel: the ACT* domains of mL-GDH$_{180}$, tyrosine hydroxylase (TH, PDB code 2MDA) (Zhang et al., 2014) and aspartate kinase (AK, PDB code 3L76) (Lang et al., 2014) as well as the ACT domain with a canonical fold of AK are superimposed (shown as ribbons). Atomic coordinates are also shown on the right in rainbow colors. The color of the labels is the same as the structures on the left. ACT* domains differ from the archetypal ACT fold in that strand β1 is located in the position usually occupied by strand β4, creating an ACT-like ββαββα topology with a $\beta_1\beta_2\beta_4\beta_3$ antiparallel sheet. Lower panel: structure based sequence alignment of the domains in the upper panel. Secondary structure elements are displayed on the alignment. Conserved core residues previously identified in the ACT family (marked with a # symbol) are shown as white positions in the structures on the right in the upper panel.

**Figure S3**

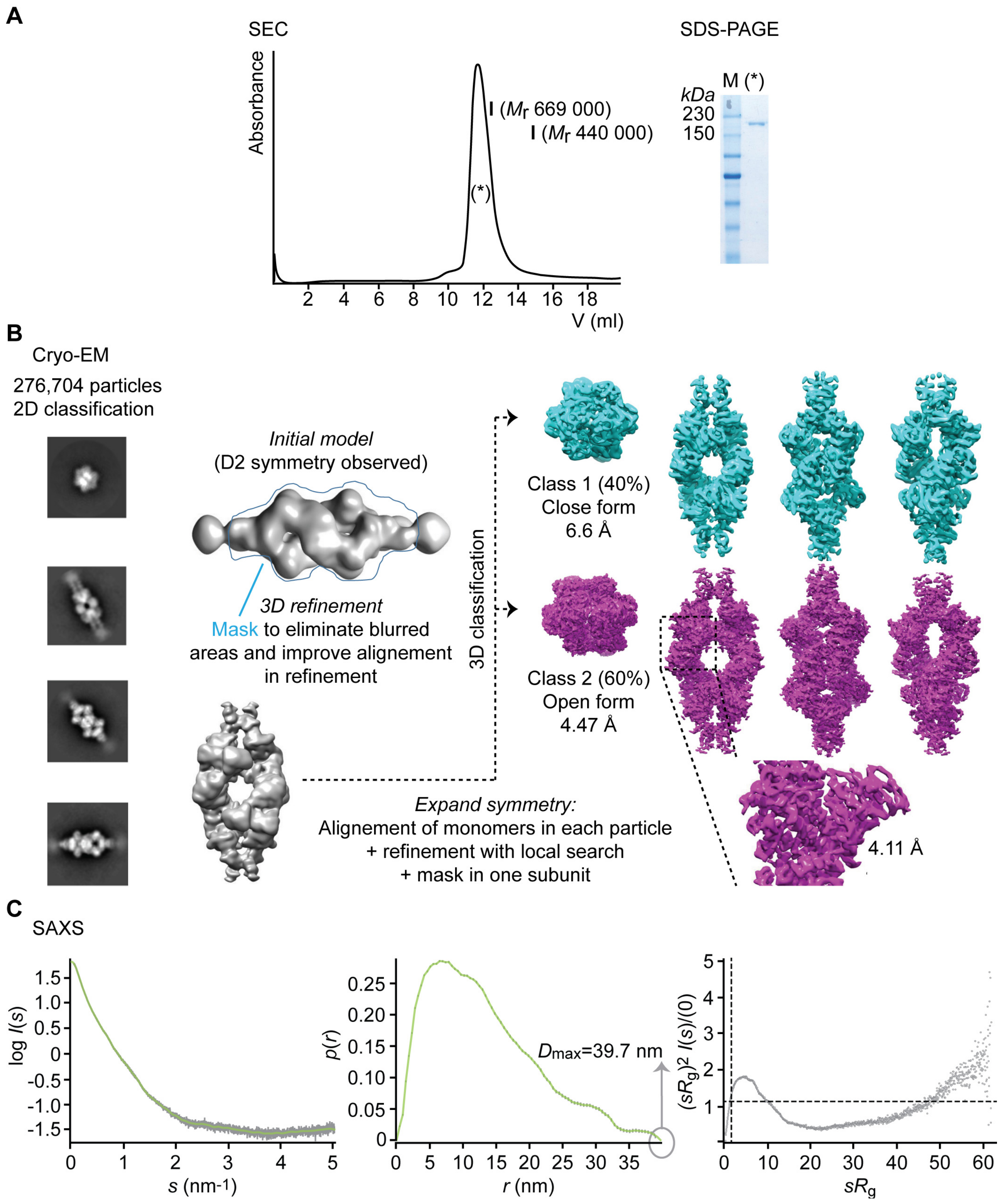


**Properties of native mL-GDH$_{180}$. (A)** Left panel: a size-exclusion chromatography using a Superose 6 10/300 GL column equilibrated in buffer 20 mM MES, 300 mM NaCl, 5 mM $MgCl_2$, pH 6.0, was performed as the final step for mL-GDH$_{180}$ purification. Thyroglobulin (669 kDa) and ferritin (440 kDa) were employed as calibration standards. $M_r$: relative molecular weight. Right panel: the purity of recombinant mL-GDH$_{180}$ (*) was evaluated by SDS-PAGE. M: molecular weight marker. Similar results were obtained for Se-Met mL-GDH$_{180}$. (**B**) Cryo-EM data of mL-GDH$_{180}$ was acquired and processed as detailed in Table 2. An initial data set of 276,704 particles was subjected to 2D and 3D class averaging in order to select the best particles. The 3D-classification of the 106,190 final particles with imposed D2 symmetry resulted in two different conformations, a close (40%) and an open form (60%), with estimated resolutions of 6.6 Å and 4.47 Å, respectively. To improve the alignment of the cryo-EM maps, we used masks to remove the blurred regions and the refinement was focused on the subunits, leading to a final resolution of 4.11 Å for a monomer in the open conformation. (**C**) SAXS data of mL-GDH$_{180}$ was acquired and processed as detailed in Table S1**.** Left panel: logarithmic plot of the scattering intensity $I(s)$ (in arbitrary units) *vs.* the momentum transfer $s$, depicting the experimental data as gray dots and the fitted curve as a green line. Center panel: pairwise distance distribution function $p(r)$ (in arbitrary units) *vs.* $r$, showing the experimental data as gray dots and the fitted curve as a green line, as for the $I(s)$ *vs.* s plot, and the estimated $D_{max}$ value. Right panel: dimensionless Kratky plot. It exhibits a non-Gaussian bell shape, consistent with a properly folded protein. Besides, the position of the maximum, far from the expected for small globular proteins, and its slow decay that does not reach zero discloses an elongated protein with flexible regions. Dashed lines indicate the position were a globular protein maximum is predicted to be located ($sR_g = 3^{1/2}$ and $(sR_g)^2\ I(s)/I(0) = 1.104$) (Bernadó, 2010; Doniach, 2001).

**Figure S4**

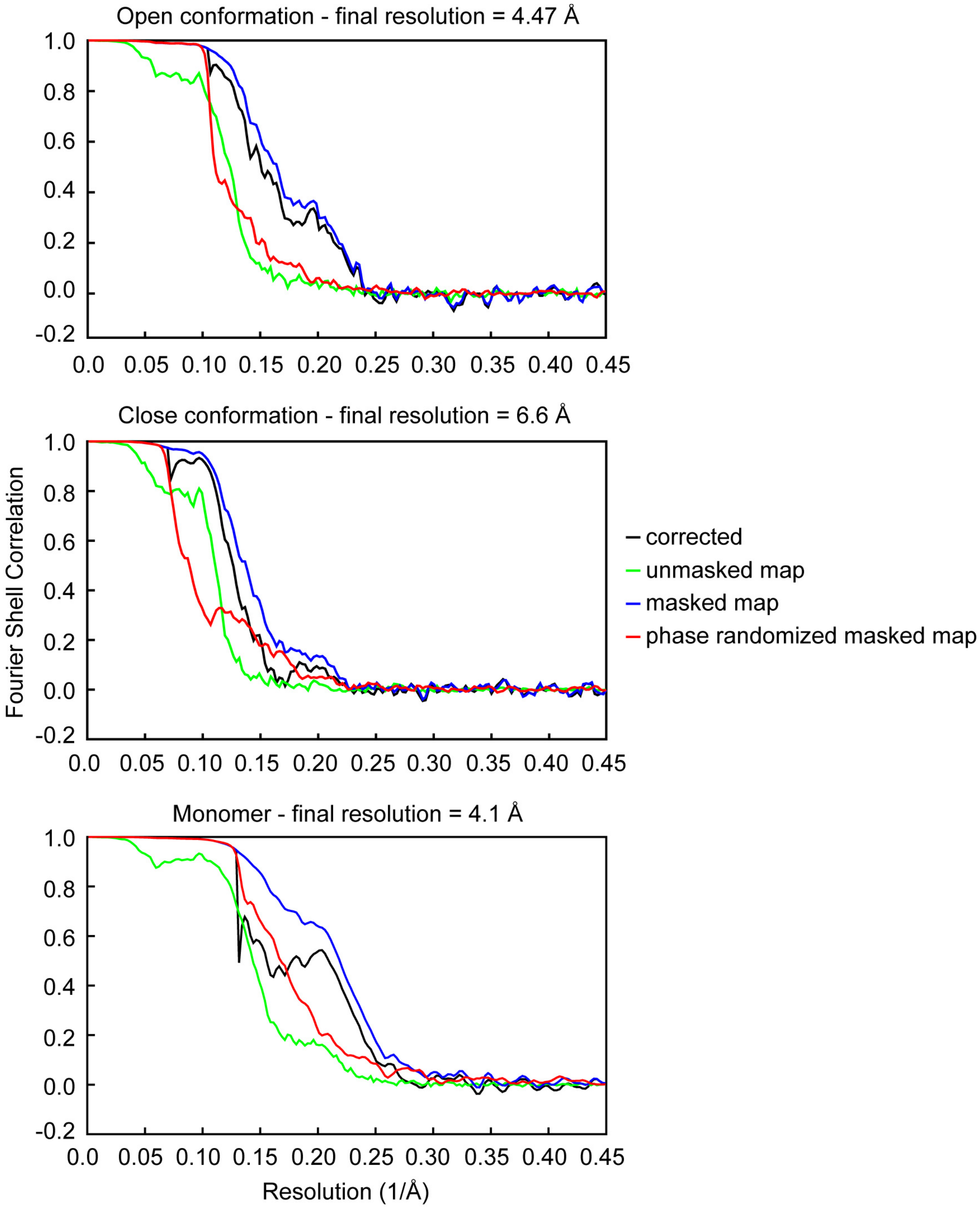


**Fourier shell correlation (FSC) for cryo-EM maps of mL-GDH$_{180}$.** The FSC is shown between independent half-maps for the three cryo-EM maps reported. Final resolutions were estimated at the 0.14 threshold of the corresponding FSC with masked density maps.

**Table S1**

**SAXS data collection and derived parameters.**

| | |
|---|---|
| **Data collection parameters** | |
| Instrument | ESRF ID14EH3 |
| Wavelength (Å) | 0.931 |
| *s* range ($Å^{-1}$)[a] | 0.009-0.6 |
| Concentration range (mg/ml) | 1.0-14.0 |
| Temperature (K) | 288 |
| **Structural parameters** | |
| $I(0)$ (relative) (from p(r)) | 64.9 ± 0.3 |
| $R_g$ (Å) (from p(r)) | 106 ± 5 |
| $I(0)$ (relative) (from Guinier) | 64.5 ± 0.3 |
| $R_g$ (Å) (from Guinier) | 101 ± 5 |
| $D_{max}$ (Å) | 39.7 |
| Porod volume estimate ($Å^3$) | 1200,000 ± 100,000 |
| Excluded volume estimate ($Å^3$) | 1300,000 ± 200,000 |
| Dry volume calculated [b] from sequence ($Å^3$) | 856,168 |
| **Molecular mass determination** | |
| Molecular mass $M_r$ (Da) (from Porod volume ($V_p$/1.7)) | 690,000 ± 70,000 |
| Molecular mass $M_r$ (Da) (from excluded volume ($V_{ex}$/2)) | 660,000 ± 80,000 |
| Calculated $M_r$ (tetramer, from sequence) | 705,071 |
| **Software employed** | |
| Data processing | PRIMUS (Franke et al., 2017; Konarev et al., 2003) |
| Porod volume calculation | DATPOROD (Franke et al., 2017) |

[a] Momentum transfer $|s| = 4\pi\sin(\theta)/\lambda$

[b] Dry volume from: http://www.basic.northwestern.edu/biotools/proteincalc.html

$M_r$: molecular mass

$R_g$: radius of gyration

$D_{max}$: maximal particle dimension

$V_p$: Porod volume

$V_{ex}$: particle excluded volume
